# Partitioning variance in a signaling trade-off under sexual selection reveals among-individual covariance in trait allocation

**DOI:** 10.1101/2022.10.10.511667

**Authors:** Michael S. Reichert, Iván de la Hera, Maria Moiron

## Abstract

Understanding the evolution of traits subject to trade-offs is challenging because phenotypes can (co)vary at both the among- and within-individual levels. Among-individual covariation indicates consistent, possibly genetic, differences in how individuals resolve the trade-off, while within-individual covariation indicates trait plasticity. There is also the potential for consistent among-individual differences in behavioral plasticity, although this has rarely been investigated. We studied the sources of (co)variance in two characteristics of an acoustic advertisement signal that trade off with one another and are under sexual selection in the gray treefrog, *Hyla chrysoscelis*: call duration and call rate. We recorded males on multiple nights calling spontaneously and in response to playbacks simulating different competition levels. Call duration, call rate, and their product, call effort, were all repeatable both within and across social contexts. Call duration and call rate covaried negatively, and the largest covariance was at the among-individual level. There was extensive plasticity in calling with changes in social competition, and we found some evidence for among-individual variance in call rate plasticity. The significant negative among-individual covariance in trait values is perpendicular to the primary direction of sexual selection in this species, indicating potential limits on the response to selection.

## INTRODUCTION

A fundamental tenet of life history theory is that variation in the allocation of resources to different phenotypic characteristics is subject to trade-offs, resulting in a negative covariance between traits (Stearns 1989; Agrawal et al. 2010; Garland 2014). Understanding how individuals resolve these trade-offs is key to understanding strategies related to important fitness-related traits such as mating, foraging decisions, growth, and parental care (Giraldeau and Caraco 2000; Badyaev and Ghalambor 2001; Simmons and Emlen 2006; Evans 2010). For trade-offs involving repeatedly expressed traits, there are two fundamental levels at which the trade-off can be expressed. First, trade-offs can be expressed at the among-individual level, in which case individuals differ consistently in the extent to which they favor one trait over the other. Second, trade-offs can be expressed at the within-individual level, where individuals adjust the allocation to each trait each time it is expressed. These sources of variation are not mutually exclusive and often there are both among- and within-individual contributions to trait variation. Nevertheless, it is important to quantify the relative contribution of among- and within-individual effects on trait covariance because these effects do not necessarily operate in the same direction (Dingemanse and Dochtermann 2013; Moiron et al. 2016; Berberi and Careau 2019), thus potentially leading to misinterpretations of patterns of covariance measured at only one level (van Noordwijk and de Jong 1986; Reznick et al. 2000; Careau and Wilson 2017). Furthermore, these sources of variation are not necessarily driven by the same evolutionary processes. Among-individual covariance potentially indicates underlying heritable covariance that could respond to or constrain correlated selection on those traits (Price and Langen 1992; Sinervo and Svensson 2002). Within-individual covariance indicates phenotypic plasticity and may reflect short-term responses to changes in the costs and benefits of different levels of trait variation (Baugh et al. 2014; Katz and Naug 2015). Although trade-offs are recognized as being critical forces in trait evolution and organismal performance, there is still mixed empirical evidence for their actual evolutionary consequences in natural systems, in part because the sources of trait covariance are rarely quantified (Roff and Fairbairn 2007; Laskowski et al. 2021; Chang et al. 2024).

Because of phenotypic plasticity in trait expression, the strength of trade-offs may be context-dependent (Sgrò and Hoffmann 2004). For instance, trade-offs are often more apparent in more challenging environmental conditions (Killen et al. 2013; Willi and Buskirk 2022). Furthermore, the relative contributions of among- and within-individual covariance on trait expression may vary across environmental conditions (Dingemanse et al. 2020). This could be the case if there is among-individual variance in phenotypic plasticity in at least one of the traits involved. In other words, some individuals may consistently make a large change in the trait expression in response to changing environmental conditions, while other individuals consistently make smaller changes or none at all (Brommer et al. 2005; Nussey et al. 2005; Araya-Ajoy and Dingemanse 2017). There is increasing attention to the causes and consequences of among-individual variance in plasticity, although empirical evidence remains limited (Nussey et al. 2007; Dingemanse et al. 2010; Dingemanse and Wolf 2013; Stamps 2016; O’Dea et al. 2022). Variation in individual plasticity could reflect underlying differences in state variables such as metabolic scope or condition (Biro and Stamps 2010; Biro et al. 2018; Dubois 2019), responsiveness to environmental stimuli (Wolf et al. 2011), personality differences (Highcock and Carter 2014), or differences in individual strategies for balancing the costs and benefits of plasticity in general (Snell-Rood 2013). If the costs and benefits of different levels of trait expression vary with environmental conditions, then these consistent among-individual differences in plasticity will also affect individual fitness, and could result in a response to selection if there is an underlying genetic component (Nussey et al. 2007).

Sexual signaling in Cope’s gray treefrog, *Hyla chrysoscelis*, is an excellent example of these complex sources of trait variation and covariation in the face of a strong trade off. Male *H. chrysoscelis* produce energetically costly advertisement calls to attract females. Females are especially attracted to calls with higher total acoustic energy (“call effort”) (Tanner et al. 2017). Call effort is allocated into two distinct traits that trade off: the duration of individual calls, and the rate at which calls are produced (Ward et al. 2013). Longer call durations and higher call rates both increase the energetic costs of signaling, and energy limitations are a likely driver of the trade-off between these two traits (Wells and Taigen 1986). Male calling is plastic with respect to conditions in the social environment. Specifically, males tend to increase call durations and decrease call rates in response to escalating competition (Ward et al. 2013). This plastic response to competition may be beneficial for mate attraction: when call efforts are held constant, there is some evidence that female *H. chrysoscelis* prefer longer calls with lower rates to shorter calls at a higher rate (Gerhardt et al. 1996), although another study found no such preference (Ward et al. 2013).

The (co)variance components of the characteristics involved in this signaling trade-off have not been quantified in *H. chrysoscelis*, and indeed some of these have received little attention in sexually selected traits in any species. First, the extent to which there are consistent among-individual differences (i.e. the repeatability) in each call characteristic has not been measured. In the closely related Eastern gray treefrog, *Hyla versicolor*, both call duration and call rate have been shown to be significantly repeatable, although the repeatability of call rate depended on year in some studies (Sullivan and Hinshaw 1992; Runkle et al. 1994; Gerhardt et al. 1996). Second, while there is evidence for within-individual covariance in call duration and call rate (Ward et al. 2013), it is unknown whether there is also a significant among-individual covariance component. This is likely to be the case because in *H. versicolor*, there was significant covariance between call duration and call period (the inverse of call rate) at both the phenotypic and genetic level (Welch et al. 2014), the latter indicating an among-individual component to the covariance. Third, while call duration and call rate are both plastic, it is unknown whether there is consistent among-individual variation in the magnitude of this plasticity in response to competition.

In this study, we investigated the role that different sources of (co)variance play in a performance trade-off that is under sexual selection by making repeated recordings in the field of male *H. chrysoscelis* both calling spontaneously and in response to playbacks simulating two different levels of competition, to address the following questions: 1. Are male call characteristics repeatable? And if so, are repeatabilities context-dependent? 2. Is there repeatable plasticity in male calling in response to a change in acoustic competition? 3. What are the relative contributions of within- and among-individual variation to the expected negative covariance between call duration and call rate?

## METHODS

We studied a population of *H. chrysoscelis* at a pond on the McPherson Preserve, Payne County, Oklahoma, USA (36.100575, -97.204080). Recordings were made during the breeding season (May – July) in 2019 and 2020, from 21:00-24:00. We searched for calling males as test subjects and removed any additional calling individual within 2 m to reduce the effects of intensive competition during the playback recordings. Male body temperature was measured at the beginning of the recording session using an infrared thermometer (Fluke 62 Max+) pointed at the body of the frog. We then made four recordings of each male using the procedure described below. Finally, each male was captured and individuals caught for the first time were marked for individual identification with a unique visible implant elastomer code on their limbs, and a photograph was taken of the distinctive black and yellow thigh pattern. Recaptured individuals were identified using these codes, with photographic recognition as an effective method to confirm the identification (Berokoff et al. 2023). After processing, all frogs were released back to the study pond. All procedures were carried out with the approval of the Institutional Animal Care and Use Committee at Oklahoma State University (protocol number AS-19-4) and with permits from the Oklahoma Department of Wildlife Conservation.

### Playback stimuli

We generated synthetic *H. chrysoscelis* calls with a custom script in R version 3.6.0 software (R Development Core Team 2021) using the seewave version 2.1.3 (Sueur et al. 2008) and tuneR version 1.3.3 (Ligges et al. 2013) packages. Call stimuli were designed to have the characteristics of a typical male *H. chrysoscelis* call, with temporal parameters designed to match the temperature of the test subject and to simulate two different levels of competition, average or high (henceforth, ‘competitive’). The call consisted of a repeated series of pulses (pulse duty cycle = 50%) with exponential rise and fall (each 50% of pulse duration). Each pulse contained two frequency bands, a low frequency peak at 1.1 kHz and a high frequency peak of 2.2 kHz that was 6 dB louder than the low frequency peak. Call pulse rates vary with temperature (Gerhardt 2005), so we generated stimuli with pulse rates that corresponded to one of four different temperatures (36.4, 49.4, 62.5, and 75.5 pulses/s at 15, 20, 25 and 30 °C, respectively). The ‘average’ stimulus had 52 pulses (linear call rise over 50 ms), and was repeated every 6 seconds. The ‘competitive’ stimulus had call characteristics of a male engaged in an especially competitive interaction (Reichert and Gerhardt 2012), with both increased number of pulses (70) and a more rapid repetition rate (one call every 4 s). Otherwise, the two stimuli were identical. Stimuli were repeated to make a playback track of 3 minutes total duration, and saved as .Wav files (44.1 kHz sampling rate). The playbacks were broadcast with an MP3 player (FiiO M3K) through a Pignose 7-100 portable speaker, at a sound pressure level of 85 dB (RMS, re: 20 µPa) at 1 m., calibrated at the beginning of the evening with a sound pressure level meter (Extech 407703A).

### Experimental design and simulated social contexts

A schematic of the experimental design is shown in Figure S1. We made recordings using digital audio recorders (Marantz PMD-660, Marantz PMD-661 or Tascam dr-100mkiii; 44.1 kHz sampling rate and 16-bit resolution in all cases) and directional microphones (Sennheiser ME-67). For each male on each night of recording, there were four consecutive recording periods corresponding to different “social contexts”. Note that the order of stimulus presentation was always the same to simulate escalating competition:

1. *Baseline recording*. Prior to playbacks we made a baseline recording of approximately 20 calls given spontaneously by the subject male.
2. *Average playback*. We next began playback of the average stimulus and recorded the male’s calling responses for the three-minute duration of the playback. We chose the average stimulus with the pulse rate most closely corresponding to the male’s body temperature (see above).
3. *Competitive playback*. We then immediately began playback of the competitive stimulus, again recording the male’s responses for three minutes, and adjusting for body temperature as above.
4. *Post-playback baseline*. Following the two playbacks we recorded another 20 spontaneous calls as a post-playback baseline.

We returned to the pond on multiple nights and continued to perform playbacks using the procedure described above. This resulted in many males being recorded multiple times on different nights in response to the same playback stimuli.

### Acoustic analyses

All advertisement calls given by males during each recording period were analyzed in Raven Pro 1.6 software (Cornell Lab of Ornithology). We measured three characteristics for each call. Call duration was measured as the difference between the onset and offset of the call. Call rate was measured on a call-by-call basis (final call of the recording excluded) as the reciprocal of the amount of time between the onset of a call and the onset of the next call. Call effort was calculated (again, excepting the final call) as the call duration multiplied by the call rate.

We note that our calculation of call rate includes the period of time in which the individual is calling, and so call rates may by definition not be independent of call duration. An alternative would be to calculate the inter-call interval as the amount of time between the offset of one call and the onset of the next, because this measure would not be automatically influenced by call duration. Nevertheless, here we focus on the trade-off between call duration and call rate, rather than inter-call interval, because only call rate has a known signaling function in this species. Receivers attend to variation in call rate, call rate variation is associated with the energetic cost of calling, and there is correlational selection on call duration and call rate (Wells and Taigen 1986; Ward et al. 2013; Tanner et al. 2017). Whether receivers attend to inter-call intervals is unknown. Furthermore, because duty cycles are relatively low in this species, call durations are substantially shorter than typical call periods, and thus there is no trade-off automatically imposed on call duration and rate due to time constraints. Indeed, in *H. versicolor*, males break the trade-off in conditions of extreme competition, resulting in positive covariance between duration and rate. To confirm that our results were not a product of our variable choice, we repeated the analyses below but using inter-call interval instead of call rate. These results are presented in the supplementary material and give the same qualitative conclusions to what we present here (Table S1-S4).

### Statistical analyses

#### Statistical model implementation

All statistical models described below were fitted using a Bayesian framework implemented in the statistical software R (v. 3.6.1, R Core Team 2019) using the R-package *MCMCglmm* (Hadfield 2010). For all models, we used parameter-expanded priors (Hadfield 2010). The number of iterations (1300000) and thinning interval were chosen for each model to ensure that the minimum MCMC effective sample sizes for all parameters were 1000. Burn-in was set to 300000 iterations. The retained effective sample sizes yielded absolute autocorrelation values lower than 0.1 and satisfied convergence criteria based on the Heidelberger and Welch convergence diagnostic. We drew inferences from the posterior modes and 95% Credible Intervals (CI). We considered fixed effects and correlations to be significant if the 95% CI did not include zero; estimates centered on zero were considered to provide support for the absence of an effect. For all models, we used a Gaussian error distribution, therefore call rate and call effort were transformed with a natural log+1 transformation prior to analyses to meet the assumption of a normal distribution.

#### Univariate analyses of repeatability of call characteristics

We performed a series of univariate general linear mixed-effects models to obtain estimates of the repeatability (R) of each call characteristic, separately for each social context. We fitted the call characteristic of interest as the dependent variable, and male identity as the random effect. As fixed effects, we included the same variables as in the multivariate model (see below). We estimated individual repeatability as the variance explained by male identity divided by the total variance. For all repeatability analyses, we included data from all males, i.e., also for those males that were only recorded once, for improved statistical power (Martin et al. 2011). Given the complexity of the multivariate models used to estimate trait covariance (see below) and the limited sample size, these univariate models applied to estimate trait repeatabilities were also used to corroborate our main findings related to trait (co)variances. Specifically, comparing repeatability estimates from the univariate and multivariate models helped to assess whether the complex multivariate models suffered from a lack of statistical power and/or poor model convergence, which reassuringly was not the case (see Results below).

Besides calculating the repeatability of each call characteristic in each social context, we also tested whether call characteristics were repeatable across the social contexts by calculating a single repeatability value for each call characteristic across all contexts. To do so, we fitted the call characteristic of interest as the dependent variable and social context as an additional fixed effect. The repeatability of each call characteristic, conditional to the variance explained by the fixed effects, was estimated as the proportion of the total phenotypic variance explained by individual variance.

#### Trait covariance among individuals across social contexts

To determine the level of covariance in the acoustic characteristics between social contexts, we fitted a four-trait multivariate general linear mixed-effects model for each of the acoustic characteristics, and investigated whether and how these call characteristics covaried between the four different social contexts. To do so, we simultaneously modelled the call characteristic (separate models for call duration, call rate and call effort) as four response variables, one for each social context (i.e., baseline, average playback, competitive playback, post-playback), and fitted an unstructured (co)variance matrix for the random effect of individual identity. In this way, we are able to estimate the covariance among individuals across contexts. The diagonal on the among-individual matrix represents the among-individual variance, which when standardized by the total phenotypic variance of each trait represents the individual repeatability. The covariances off the diagonal of the matrix indicate whether there are differences among individuals in plastic responses to social context (interpreted through correlations and/or changes in variances in the same matrix). The interpretation of individual variation in plastic responses through correlations is possible by assessing whether the 95% CI of the cross-context correlation includes r = +1 or not. When the 95% CI of the cross-context correlation includes +1, it means that the among-individual variance is the same in both environments (i.e., variance across contexts is constant and there is a perfect correlation of individual variation expressed in both environments) and consequently implies an absence of individual-by-environment (hereafter, IxE) interactions (i.e., no individual variance in plasticity). When the 95% CI of the cross-context correlation is different from +1, it indicates that the among-individual variance changes between the two environments (i.e., no constant variance across contexts and hence the correlation falls lower than the perfect +1), ultimately providing evidence for IxE interactions (i.e. individuals vary in plasticity). We also modeled the covariance at the within-individual level by fitting an unstructured (co)variance matrix on the residual components to control for repeated measures of calling traits on each night of sampling (Mitchell and Houslay 2021). There was no evidence for within-individual cross-context covariance for any of the three call characteristics (Table S5).

For each multivariate model, we fitted as fixed effects year (categorical variable, two levels: 2019 and 2020), time of night (continuous variable, in hours past 21:00), calendar day (continuous variable), and male body temperature (continuous variable). Both time of night and calendar day were centered, by subtracting the mean, and standardized, by dividing by the standard deviation. Additionally, we fitted a “sequence” effect, a continuous variable numbering the calls in order on the recording, to control for potential autocorrelation among the repeated observations.

#### Sources of covariance in the trade-off between call duration and call rate across social contexts

We fitted bivariate general linear mixed-effects models to investigate whether and how call duration and call rate covaried across levels (at the population, among-individual and within-individual levels) in two different social contexts (separate models for the average and competitive playbacks). To do so, we simultaneously modelled call duration and call rate as response variables, and fitted unstructured (co)variance matrices for the random effects of individual identity and residual variance. We fitted the same fixed effects as for the four-trait multivariate model (see above).

## RESULTS

The dataset includes 38 males recorded on more than one night (2 nights: N=24, 3 nights: N=5, 4 nights: N=6, 5 nights: N=3). In addition, data from 77 males that were recorded on only one night are included. Raw data and analysis code are deposited in Dryad: https://doi.org/doi.10.5061/dryad.kd51c5b95.

### Repeatability of call characteristics

All male call characteristics were significantly repeatable in each of the four social contexts (Figure 1). Social context did not generally affect the repeatability coefficients, which were largely similar across all four contexts (Table S6).

When we calculated a single repeatability value for each call characteristic across all social contexts, we found again that all call characteristics were significantly repeatable (Table S6). The magnitudes of the repeatability coefficients when pooling together data from the four contexts were approximately the same as the averaged repeatability across the four contexts (Figure 1). Finally, as expected, the individual repeatabilities estimated from the multivariate model of calling across contexts, which were extracted from the 4x4 matrix and calculated as the trait variance in each context divided by the total variance, resembled the repeatability estimates using univariate models (Table 1 vs. Table S6, respectively).

Fixed effects for the four-trait multivariate models for each call characteristic are given in Table S7. There was an effect of temperature on call duration and call rate across all social contexts. At higher temperatures, males had lower call durations, likely due to increased pulse repetition rates (Gayou 1984), and called at a faster rate. Call effort in the baseline and competitive playback was also influenced by temperature, with lower calling effort at higher temperatures. Calendar day also almost always had an effect on all call characteristics across the four contexts: males had shorter calls, and lower call efforts and call rates later in the season. Call characteristics differed between years of sampling, with higher call rates but lower call durations and effort in year 2020 compared to year 2019. Finally, time of the night measured as hours after 21h00 only had a significant negative effect on call duration, but not on rate or effort. Males had shorter calls later in the night than earlier.

**Figure 1.**
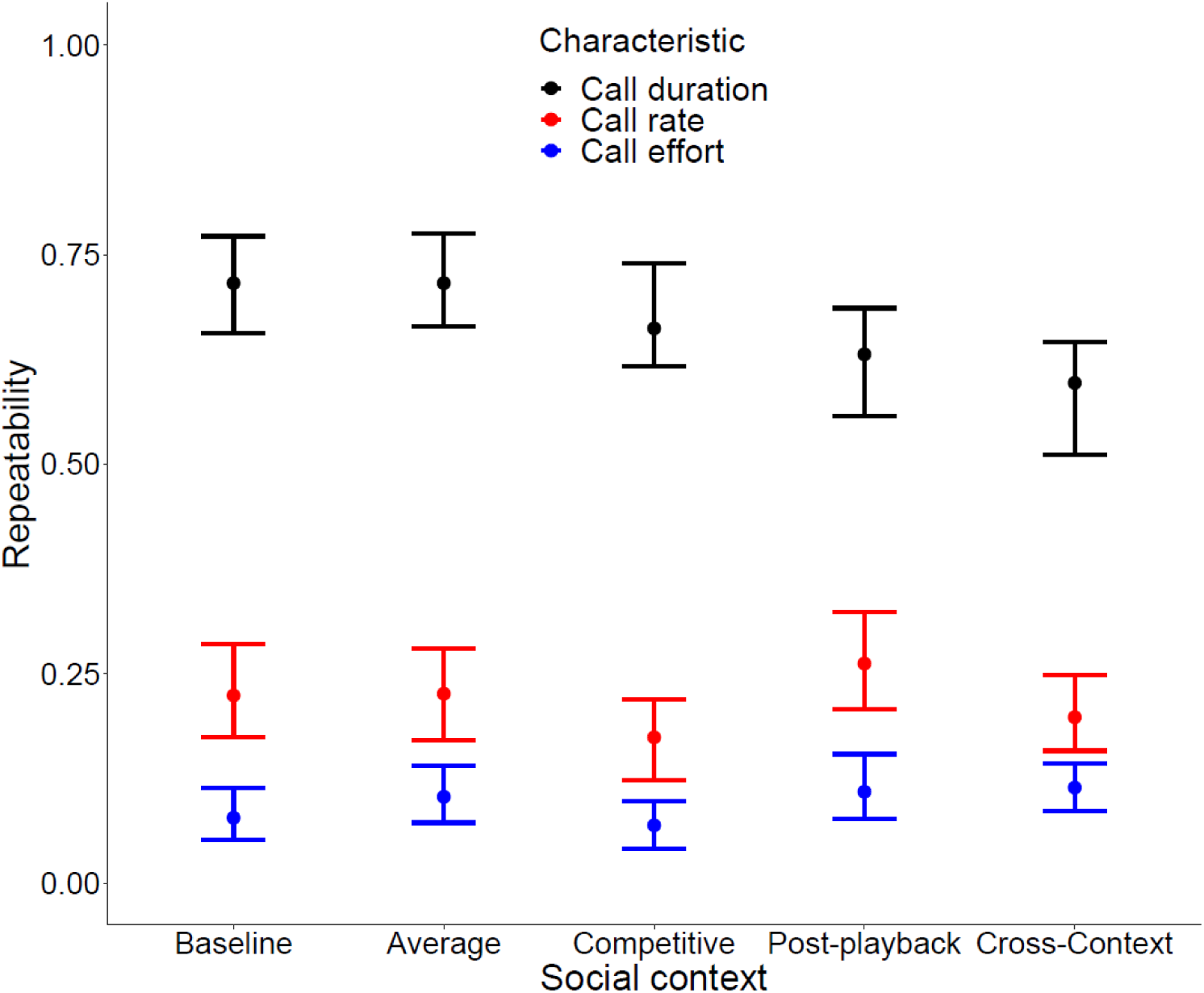
Repeatability estimates of call characteristics in each recording period (social context) and the composite ‘cross-context’ repeatability calculated from data from all social contexts (Table S6). Repeatability estimates are shown as posterior modes with 95% Credible Intervals, calculated from univariate models of the given call characteristic in a single social context. Colors indicate the different call characteristics.

**Table 1.**
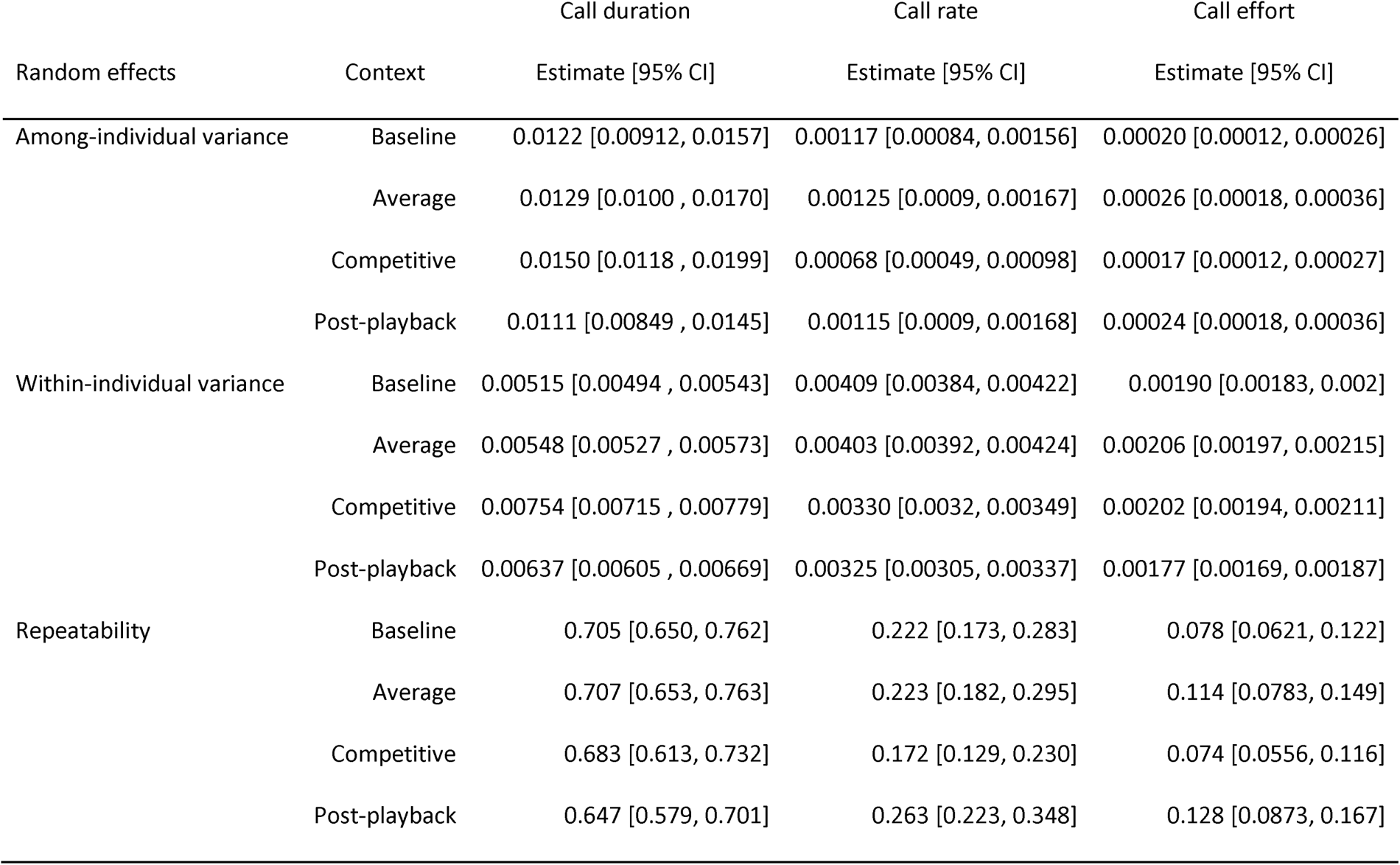
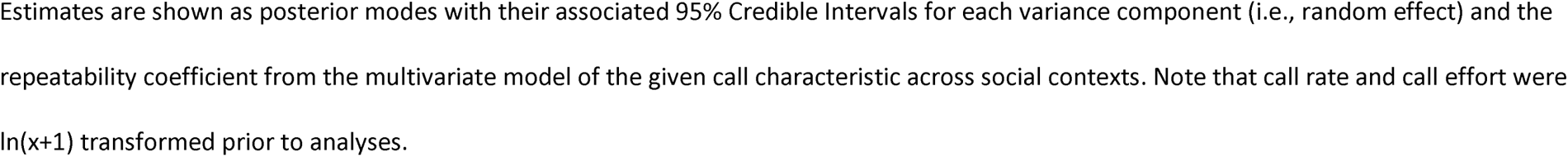
Sources of phenotypic variance in call characteristics across social contexts.

### Individual differences in call characteristic plasticity

Call rate showed modest evidence for individual variation in plasticity across social contexts, as indicated by the different-from-one correlations across contexts (Table 2). In other words, there was among-individual variation in how males changed their call rates in response to changes in the simulated level of competition. Although these correlations differed from one (i.e., the upper 95% CI was not one), they were very strong in all cases, ranging from 0.822 to 0.919. This individual variation in plasticity was particularly clear for the change in call rate between the baseline to the average playback (Figure 2A, Table 2). In line with this finding based on the assessment of cross-context correlations, when we assessed the levels of among-individual variance within each social context, we found that there was slightly less variation in call rate in the competitive playback context compared to the other three social contexts, and that the 95% CI were only slightly overlapping between variance estimates (Table 1). This pattern again is modest evidence for IxE in call rate. If individuals did not differ in how they responded to increased social competition, then they should have all behaved in a similar fashion across contexts and therefore the among-individual variance should not have changed (i.e. patterns of response would have been parallel across individuals). For call duration (Table 2) and call effort (Figure 2C; Table 2), however, we did not find clear evidence for differences in the plastic responses to any of the four changes in context, as indicated by the close to one cross-context correlations and the similar among-individual variances. These results suggest that there was no significant variation among individuals in the extent to which they changed call duration or call effort as social competition increased. A possible exception is for call duration in the baseline and competitive contexts (Figure 2B), where the correlation differs from one and is of a similar magnitude to those found for call rate (Table 2), which may indicate IxE for call duration for these contexts.

**Figure 2.**
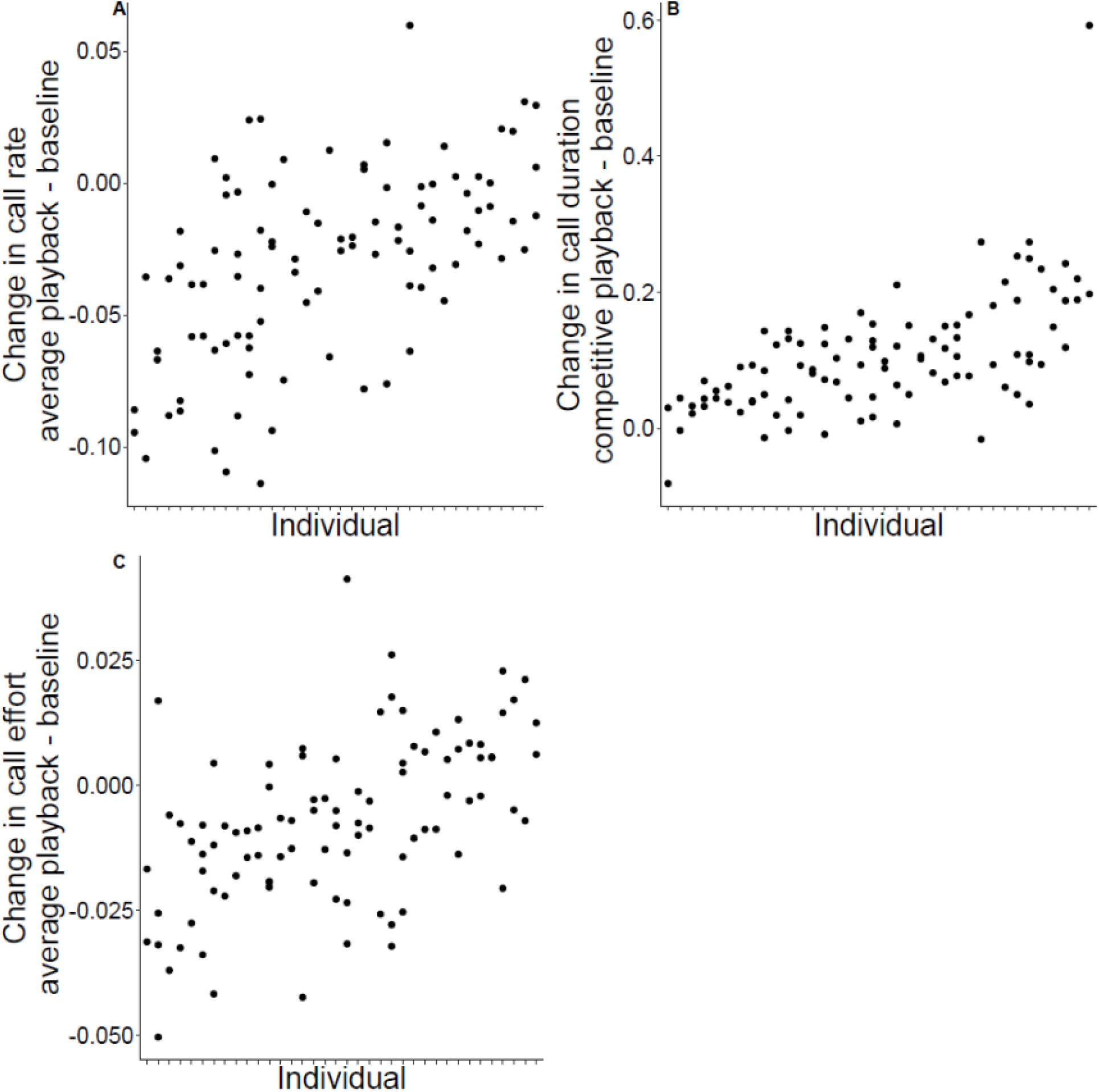
Illustration of among-individual variation in calling plasticity. Points are arranged by individual identity on the x-axis (thus points on the same location on the x-axis correspond to recordings of the same individual taken on different nights), with individuals with the smallest mean value of the trait on the y-axis at the left, and individuals with the largest mean value of that trait on the right, and the y-axis gives the difference between the call characteristic measured during the playback (average playback stimulus for call rate and call effort, competitive playback stimulus for call duration) and the same call characteristic measured during the baseline. Only individuals with data from two or more nights are included. A. Change in mean call rate (calls/s) between average playback and baseline. B. Change in mean call duration (s) between competitive playback and baseline. C. Change in mean call effort between average playback and baseline. For call rate and call duration in these contexts, there was evidence of IxE interactions, while there was no evidence of this for call effort. Individual changes in call characteristics were statistically modeled with a cross-context bivariate model, but they are represented here as raw differences for illustrative purposes only.

**Table 2.** Among-individual estimates for covariance and correlations in call characteristics across social contexts.

| Contexts compared | Call duration | Correlation<br>[95% CI] | Call rate | Correlation<br>[95% CI] | Call effort | Correlation<br>[95% CI] |
| --- | --- | --- | --- | --- | --- | --- |
|  | Covariance [95%<br>CI] |  | Covariance [95% CI] |  | Covariance [95% CI] |  |
| Average – baseline | 0.0108 [0.00864,<br>0.0150] | 0.916 [0.870,<br>0.942] | 0.00098 [0.00068,<br>0.0013] | 0.822 [0.691,<br>0.884] | 0.00018 [0.00014,<br>0.00027] | 0.950 [0.813,<br>0.984] |
| Competitive –<br>baseline | 0.0104 [0.00809,<br>0.0147] | 0.796 [0.740,<br>0.879] | 0.00070 [0.00051,<br>0.001] | 0.826 [0.693,<br>0.893] | 0.00018 [0.00013,<br>0.00024] | 0.983 [0.913,<br>0.998] |
| Competitive –<br>average | 0.0135 [0.00973,<br>0.0168] | 0.935 [0.885,<br>0.953] | 0.00086 [0.00061,<br>0.00116] | 0.919 [0.837,<br>0.963] | 0.00019 [0.00014,<br>0.00027] | 0.931 [0.848,<br>0.996] |
| Post-playback –<br>baseline | 0.0109 [0.00808,<br>0.0140] | 0.912 [0.861,<br>0.942] | 0.00096 [0.00075,<br>0.00141] | 0.867 [0.769,<br>0.926] | 0.00020 [0.00016,<br>0.00029] | 0.980 [0.896,<br>0.997] |
| Post-playback –<br>average | 0.0112 [0.0087,<br>0.0149] | 0.941 [0.911,<br>0.965] | 0.00096 [0.00071,<br>0.00139] | 0.824 [0.743,<br>0.905] | 0.00022 [0.00016,<br>0.00032] | 0.921 [0.806,<br>0.986] |
| Post-playback -<br>competitive | 0.0113 [0.00896,<br>0.0155] | 0.901 [0.868,<br>0.941] | 0.00075 [0.00055,<br>0.00109] | 0.853 [0.756,<br>0.927] | 0.00020 [0.00015,<br>0.00029] | 0.972 [0.892,<br>0.997] |

Estimates show the posterior mode with its associated 95% Credible Intervals for the covariance and correlation at the among-individual level
for each call characteristic across each pairwise combination of social contexts. Cross-context correlation coefficients that are clearly different
from one provide evidence for among-individual variance in calling plasticity, in other words that there is an IxE interaction.

### Sources of covariance in the trade-off between call duration and call rate across social contexts

For the average playback, there was a strong trade-off between call duration and call rate; longer duration calls were given at lower rates at the phenotypic level (Table 3). When we tested whether this phenotypic negative correlation was also present at other hierarchical levels of variation, we observed that there was a significant negative covariance between call duration and call rate both within-(residual variance) and among-males (Table 3). The negative correlation among individuals indicates that some males consistently resolved the trade-off by producing longer-duration calls at a lower rate than others. The negative within-individual correlation suggests that on a call-to-call basis, when a male had a high call duration it also had a low call rate. Across the three levels of correlations, the among-individual correlation was the strongest, the phenotypic correlation was intermediate, and the within-individual correlation was the weakest (Table 3).

When we tested for the existence of a trade-off between call duration and call rate for the competitive playback context, the pattern was very similar to what we observed for the average playback context. Specifically, we also found a negative correlation between these two call traits at the phenotypic-, among- and within-individual level, although the within-individual correlation was not significant. This finding indicates that some males also resolved the trade-off by producing on average longer-duration calls at a lower rate than others during the most competitive social context, but on a call-to-call basis individuals were not adjusting their call rate based on the duration of that specific call (Table 3). Fixed effects on both bivariate models are shown in Table S8.

**Table 3.**
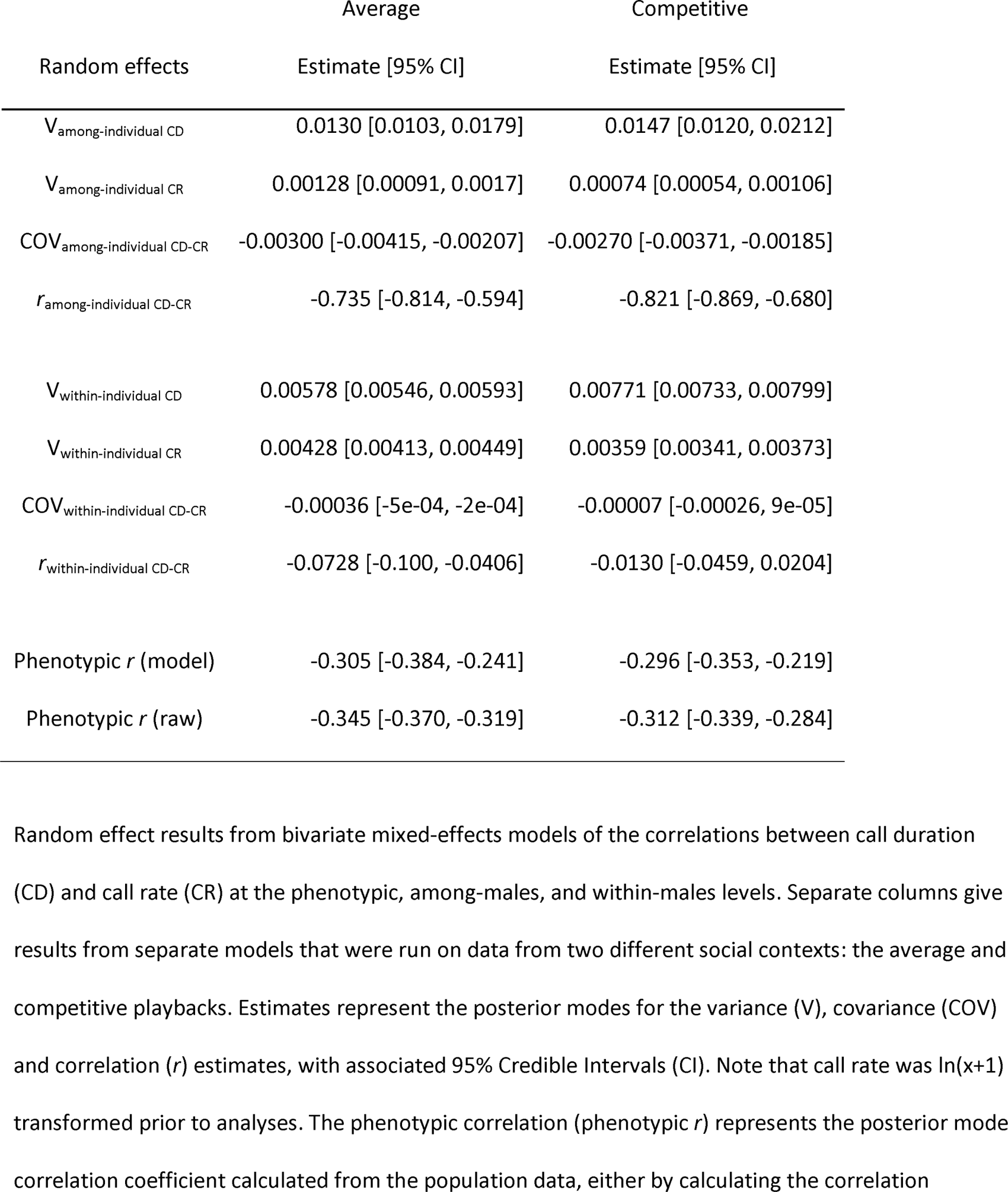
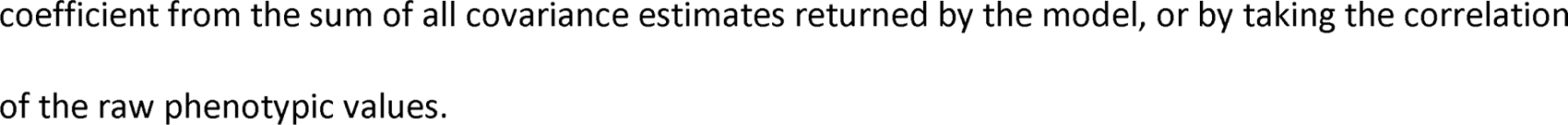
Results from a bivariate model testing for the covariance between call duration (CD) and call rate (CR)

## DISCUSSION

We found that both phenotypic plasticity in response to competition and repeatable among-individual differences contributed to variation in call characteristics in H. chrysoscelis. Furthermore, for two key temporal call characteristics, call duration and call rate, there was significant negative covariation, confirming the expected trade-off between these two characteristics at the phenotypic level (Ward et al. 2013). Because we had multiple recordings per individual, we could partition the covariance into its components at different levels of phenotypic variation. We found that the among-individual covariance between call duration and call rate was greater than the within-individual covariance. Thus, although males exhibited short-term behavioral plasticity by adjusting their calls in accordance with the trade-off between duration and rate in response to competition, some individuals nevertheless consistently allocated more calling energy to call duration (and less to call rate) than others. Finally, there was some evidence for consistent among-individual differences in the plastic response to competition, in particular for call rate, where some males consistently changed their call rates more than others when exposed to an increase in competition. We discuss below the implications and limitations of these results for the understanding of sexual selection and of the evolution of multivariate phenotypes subject to trade-offs.

### Implications for sexual selection

Call duration was highly repeatable both within and across social contexts representing different levels of competition. The cross-context repeatability indicates that even though males vary their call duration in response to competition, those males that produce the longest duration calls in low levels of competition are usually the same males that produce the longest duration calls in high levels of competition. Thus, males should maintain more or less the same rank order of call duration as levels of competition within the chorus fluctuate, as has also been noted in H. versicolor (Schwartz et al. 2002). Call rate and call effort were also repeatable, although the magnitude of the repeatability coefficient was much smaller for these characteristics, indicating a greater contribution of within-individual variance. The lower repeatability for call rate than call duration indicates that despite the general correlation between the two characteristics, call rate maintains enough independent variance that the within-individual variance component for call rate predominates. Call effort is the product of call rate and call duration, so the relatively low repeatability of call rate may partially explain why call effort had a low repeatability. Furthermore, the strong upper limit on call effort (Reichert and Gerhardt 2012) may restrict the among-male variance for this characteristic. The fact that call duration shows relatively greater among-male variance may explain why call duration often outweighs call rate in female mate choice decisions (Gerhardt et al. 1996). Females sample male calls over a limited time period (Schwartz et al. 2004), thus it makes sense that call duration, which has very high repeatability and likely can be reliably assessed by attending to just a few calls, predominates over call rate, which is much less repeatable and probably more difficult to assess (Tanner and Bee 2020).

We found that, in response to playbacks of calls with increased duration, males typically increased their own call durations while reducing call rates (Figure 2). This finding corroborates the results of previous studies (Ward et al. 2013) and indicates that the trade-off is resolved in favor of call duration at higher levels of competition. However, it remains unclear why the trade-off is resolved in this direction, and whether this is driven by competition among males or to attract mate searching females, or both. Males certainly attend to competitors and their behavior in our playback trials, indicating that they are capable of detecting differences in these call characteristics (Figure 2B). Nevertheless, whether longer calls are actually more effective at repelling rival males is unknown. If this is the case, then the repeatability of call duration and the strong among-individual component of the covariance between call duration and call rate would result in some males being consistently more competitive than others. In terms of female preferences, there is mixed evidence that call duration predominates over call rate in female decisions (Gerhardt et al. 1996; Ward et al. 2013), and preference strength for either characteristic weakens as its magnitude increases, which should weaken selection towards either extreme of the trade-off axis (Tanner et al. 2017). There may be other advantages for males to resolve the trade-off by producing longer calls at lower rates, for instance if longer calls could stand out better against the background noise of the chorus, although evidence for such a benefit is limited (Bee and Schwartz 2013).

### Individual variation in plasticity

As for variation in individual traits, repeatedly measuring plasticity in response to changing environmental contexts and then partitioning the variance in plasticity into its among- and within-individual components gives important insights into the causes and evolutionary consequences of limited plasticity (Dingemanse et al. 2010; Stamps and Biro 2016). There is accumulating evidence that behavioral plasticity can be repeatable (Mathot et al. 2011; Morand-Ferron et al. 2011; Mitchell and Biro 2017). If this repeatability indeed reflects genetic variation, then there is potential for selection to act on variation in individual plasticity (Roff 1997; Nussey et al. 2007; Dingemanse et al. 2012; Brommer 2013). Among-individual variance in plasticity could also affect the strength of trade-offs in different environments (Sgrò and Hoffmann 2004). We found some evidence for consistent among-individual differences in behavioral plasticity for call rate; in response to changes in the level of social competition some males consistently changed their call rates more than others. The ability to respond to a competitor through plastic changes in calls likely affects success in mate competition for male H. chrysoscelis, and our findings indicate that for call rate there is a moderate potential for a response to selection on calling plasticity. However, for call duration and call effort there was little evidence for repeatable individual plasticity. As a result, the overall covariance between call duration and call rate and the contributions of among- and within-individual covariance did not strongly differ across contexts.

Given that we found that call duration both was repeatable and plastic with respect to the level of competition, we expected that plasticity in call duration in response to competition would also be repeatable but there was only evidence for this for the comparison of calling in the baseline and in response to the competitive playback. It is unclear why there was not stronger evidence for repeatable variation in call duration plasticity. One reason may be that we measured male responses in the field, and therefore social conditions were not standardized. On any given night, males may have experienced different levels of competition, and both prior experience and current levels of competition are known to affect male frogs’ responses to competitors (Humfeld et al. 2009; Reichert 2010; Heap et al. 2012). We attempted to remove the nearest competitors prior to recordings, but there was certainly still variation in background calling levels that could not be controlled for, and this could have introduced noise in our measurements. Although recording males in standardized conditions (e.g. a quiet laboratory chamber) may have given us greater power to detect among-individual variation in male responses to competition, this is not the biologically-relevant environment in which sexual selection is acting. Another potential explanation of why we did not detect individual differences in plasticity is limited statistical power. Detecting repeatable plasticity in multivariate models requires large sample sizes (Martin et al. 2011; van de Pol 2012), and it is possible that our sample was not sufficient for this purpose. However, several of our results suggested that our model was not underpowered. First, although some among-individual variance components were small there were no identified issues with model convergence and the posterior distribution of these values did not converge towards zero. Second, the repeatability coefficients calculated from the multivariate model were very similar in magnitude and variation to those calculated from univariate models (Table 1 vs Table S6). Because univariate models with a simpler random effect model structure are in principle much less likely to be underpowered, the close correspondence between univariate and multivariate repeatability estimates suggests that the multivariate model had sufficient power to estimate variance components.

We examined plasticity at the level of short-term variation in response to an immediate change in social competition. Males may also plastically adjust their calls at other temporal scales. For instance, we found that call characteristics often varied with both time of night and calendar day, typically leading to reduced energetic expenditures later at night and later in the year. Similarly, in H. versicolor, males spent less time calling per night as the season progressed, although their actual call efforts while calling did not change (Runkle et al. 1994). Our sampling does not allow us to partition the sources of variance in calling at these longer temporal scales, but it is conceivable that males differ in the extent to which they allocate energy to calling within a night or across the breeding season. Given the importance of attendance at the breeding chorus to mating success, among-individual variance in calling plasticity at these temporal scales could have important consequences for sexual selection. Furthermore, this could lead to temporal variation in the resolution to the trade-off between call duration and call rate.

### Trade-offs and sources of covariance

A negative phenotypic correlation between traits is the hallmark of a trade-off, but these data alone are insufficient to determine at what level(s) of variation the trade-off takes place, and therefore what the consequences are for behavioral expression and trait evolution. Indeed, a negative covariance between traits measured at the population level may have zero or even positive covariance for specific variance components. Therefore, in the absence of pedigreed or genetic data, it is necessary to take repeated measurements of all traits involved in a trade-off to partition among the different levels of covariance (Dingemanse and Dochtermann 2013; Moiron et al. 2016; Berberi and Careau 2019). In our case, we found negative covariance between call duration and call rate at every level of variation. This likely reflects the fact that the high energetic costs of increasing both call duration and call rate are applicable to both within- and among-individual variation, ultimately imposing a trade-off at both levels. Nevertheless, we found that the covariance was the strongest at the among-individual level, and was much weaker at the level of within-male variation from one call to the next. One caveat is that because we recorded males in natural choruses, some among-individual covariance could be environmental in origin, if males consistently differ in their tendency to call from more or less competitive areas of the pond (Aplin et al. 2015). We controlled for some social factors in our models, but until genetic covariances are measured in this species we cannot rule out that environmental sources could have inflated estimates of among-individual covariance (Niemelä and Dingemanse 2017).

In the closely-related H. versicolor, there was a significant genetic correlation between call duration and call rate that was of a similar magnitude to their phenotypic correlation (Welch et al. 2014). Our finding of significant among-individual phenotypic covariance between call duration and call rate in H. chrysoscelis therefore indicates the potential for genetic covariance in this species as well. The evolutionary significance of a negative among-individual covariance is that, if this indeed has a genetic underpinning, there is potential for a correlated response to selection (Lande and Arnold 1983; Blows and Hoffmann 2005; Walsh and Blows 2009). However, the strongest female preferences in this species are for calls with both longer call durations and higher call rates, and thus higher call efforts (Tanner et al. 2017), and we found that call effort was only weakly repeatable. Thus, the response to selection on call characteristics may be limited, because most genetic variance would be perpendicular to the major axis of sexual selection (McGuigan et al. 2008). One final consideration is that the male advertisement call is an integrated phenotype made up of many components (Reichert and Höbel 2018), and our analyses focused on only two of these. We did not examine a third characteristic, call amplitude, which is also under sexual selection (Fellers 1979) and contributes to the energetic cost of calling (Prestwich 1994). Although within-individual variation in call amplitude appears to be minimal (Gerhardt 1975; Love and Bee 2010), whether there are consistent among-individual differences in amplitude, and whether it covaries with other call traits, is unknown. An important goal for future studies is therefore to attempt to characterize more fully the interrelationships among many potentially covarying traits, and determine at what levels they (co)vary (Blows and Hoffmann 2005; Walsh and Blows 2009).

## AUTHOR CONTRIBUTIONS

M.S.R. conceived the study, M.S.R and I.d.H collected data, M.M. and M.S.R. analyzed data, M.S.R. wrote the manuscript with contributions from M.M. and I.d.H.

## CONFLICT OF INTEREST STATEMENT

The authors declare no conflict of interest

## ACKNOWLEDGEMENTS

Nicole Clapp, James Erdmann, A.J. Hager and Cheyenne Smith assisted with the field work. Rachel Atherton contributed to the acoustic analyses. M.M. was funded by an Alexander von Humboldt Research Fellowship for Postdoctoral Researchers.

## DATA ACCESSIBILITY

Raw data and analysis code are deposited in Dryad: https://doi.org/doi.10.5061/dryad.kd51c5b95

## Notes

https://doi.org/doi:10.5061/dryad.kd51c5b95

### Competing Interest Statement

The authors have declared no competing interest.

### Summary of Updates

Some additional analyses and minor clarifications to the text.

## REFERENCES

Agrawal, A. A., J. K. Conner, and S. Rasmann. 2010. Tradeoffs and negative correlations in evolutionary ecology. Pp. 243–268 in M. Bell, W. Eanes, D. Futuyma, and J. Levinton, eds. Evolution after Darwin: The first 150 years. Sinauer, Sunderland, MA.

Aplin, L. M., J. A. Firth, D. R. Farine, B. Voelkl, R. A. Crates, A. Culina, C. J. Garroway, C. A. Hinde, L. R. Kidd, I. Psorakis, N. D. Milligan, R. Radersma, B. L. Verhelst, and B. C. Sheldon. 2015. Consistent individual differences in the social phenotypes of wild great tits, Parus major. Anim. Behav. 108:117–127.

Araya-Ajoy, Y. G., and N. J. Dingemanse. 2017. Repeatability, heritability, and age-dependence of seasonal plasticity in aggressiveness in a wild passerine bird. J. Anim. Ecol. 86:227–238.

Badyaev, A. V., and C. K. Ghalambor. 2001. Evolution of life histories along elevational gradients: Trade-off between parental care and fecundity. Ecology 82:2948–2960.

Baugh, A. T., K. Van Oers, N. J. Dingemanse, and M. Hau. 2014. Baseline and stress-induced glucocorticoid concentrations are not repeatable but covary within individual great tits (Parus major). Gen. Comp. Endocrinol. 208:154–163.

Bee, M. A., and J. J. Schwartz. 2013. Calling in gray treefrog choruses: modifications and mysteries. Proc. Meet. Acoust. 19:010054–010054.

Berberi, I., and V. Careau. 2019. Performance trade-offs in wild mice. Oecologia 191:11–23.

Berokoff, J., I. de la Hera, and M. S. Reichert. 2023. Image processing of thigh color pattern is an effective method for identifying individual Cope’s gray treefrogs, Hyla chrysoscelis. Ichthyol. Herpetol. 111:612–620.

Biro, P. A., and J. A. Stamps. 2010. Do consistent individual differences in metabolic rate promote consistent individual differences in behavior? Trends Ecol. Evol. 25:653–659.

Biro, P., T. Garland, C. Beckmann, B. Ujvari, F. Thomas, and J. Post. 2018. Metabolic scope as a proximate constraint on individual behavioral variation: effects on personality, plasticity, and predictability. Am. Nat. 192:142–154.

Blows, M. W., and A. A. Hoffmann. 2005. A reassessment of genetic limits to evolutionary change. Ecology 86:1371–1384.

Brommer, J. E. 2013. Phenotypic plasticity of labile traits in the wild. Curr. Zool. 59:485–505.

Brommer, J. E., J. Merilä, B. C. Sheldon, and L. Gustafsson. 2005. Natural selection and genetic variation for reproductive reaction norms in a wild bird population. Evolution 59:1362–1371.

Careau, V., and R. S. Wilson. 2017. Of uberfleas and krakens: Detecting trade-offs using mixed models. Integr. Comp. Biol. 57:362–371.

Chang, C., M. Moiron, A. Sánchez-Tójar, P. Niemelä, and K. Laskowski. 2024. What’s the meta-analytic evidence for life-history trade-offs at the genetic level? Ecol. Lett.

Dingemanse, N. J., I. Barber, and N. A. Dochtermann. 2020. Non-consumptive effects of predation: does perceived risk strengthen the genetic integration of behaviour and morphology in stickleback? Ecol. Lett. 23:107–118.

Dingemanse, N. J., I. Barber, J. Wright, and J. E. Brommer. 2012. Quantitative genetics of behavioural reaction norms: genetic correlations between personality and behavioural plasticity vary across stickleback populations. J. Evol. Biol. 25:485–496.

Dingemanse, N. J., and N. A. Dochtermann. 2013. Quantifying individual variation in behaviour: mixed-effect modelling approaches. J. Anim. Ecol. 82:39–54.

Dingemanse, N. J., A. J. N. Kazem, D. Réale, and J. Wright. 2010. Behavioural reaction norms: animal personality meets individual plasticity. Trends Ecol. Evol. 25:81–89.

Dingemanse, N. J., and M. Wolf. 2013. Between-individual differences in behavioural plasticity within populations: causes and consequences. Anim. Behav. 85:1031–1039.

Dubois, F. 2019. Why are some personalities less plastic? Proc. R. Soc. B Biol. Sci. 286:20191323.

Evans, J. P. 2010. Quantitative genetic evidence that males trade attractiveness for ejaculate quality in guppies. Proc. R. Soc. B Biol. Sci. 277:3195–3201.

Fellers, G. M. 1979. Aggression, territoriality, and mating behaviour in North American treefrogs. Anim. Behav. 27:107–119.

Garland, T. 2014. Trade-offs. Curr. Biol. 24:R60–R61.

Gayou, D. C. 1984. Effects of temperature on the mating call of Hyla versicolor. Copeia 1984:733–738.

Gerhardt, H. C. 2005. Acoustic spectral preferences in two cryptic species of grey treefrogs: implications for mate choice and sensory mechanisms. Anim. Behav. 70:39–48.

Gerhardt, H. C. 1975. Sound pressure levels and radiation patterns of the vocalizations of some North American frogs and toads. J. Comp. Physiol. A 102:1–12.

Gerhardt, H. C., M. L. Dyson, and S. D. Tanner. 1996. Dynamic properties of the advertisement calls of gray tree frogs: patterns of variability and female choice. Behav. Ecol. 7:7–18.

Giraldeau, L. A., and T. Caraco. 2000. Social Foraging Theory. Princeton University Press, Princeton.

Hadfield, J. D. 2010. MCMCglmm: MCMC methods for multi-response GLMMs in R. J. Stat. Softw. 33:1–22.

Heap, S., D. Stuart-Fox, and P. Byrne. 2012. Variation in the effect of repeated intrusions on calling behavior in a territorial toadlet. Behav. Ecol. 23:93–100.

Highcock, L., and A. J. Carter. 2014. Intraindividual variability of boldness is repeatable across contexts in a wild lizard. PLoS One 9:e95179.

Humfeld, S. C., V. T. Marshall, and M. A. Bee. 2009. Context-dependent plasticity of aggressive signalling in a dynamic social environment. Anim. Behav. 78:915–924.

Katz, K., and D. Naug. 2015. Energetic state regulates the exploration-exploitation trade-off in honeybees. Behav. Ecol. 26:1045–1050.

Killen, S. S., S. Marras, N. B. Metcalfe, D. J. McKenzie, and P. Domenici. 2013. Environmental stressors alter relationships between physiology and behaviour. Trends Ecol. Evol. 28:651–658. Elsevier Ltd.

Lande, R., and S. J. Arnold. 1983. The measurement of selection on correlated characters. Evolution 37:1210–1226.

Laskowski, K. L., M. Moiron, and P. T. Niemelä. 2021. Integrating behavior in life-history theory: Allocation versus acquisition? Trends Ecol. Evol. 36:132–138.

Ligges, U., S. Krey, O. Mersmann, and S. Schnackenberg. 2013. tuneR: Analysis of music and speech. URL: https://CRAN.R-project.org/package=tuneR.

Love, E. K., and M. A. Bee. 2010. An experimental test of noise-dependent voice amplitude regulation in Cope’s grey treefrog, Hyla chrysoscelis. Anim. Behav. 80:509–515.

Martin, J. G. A., D. H. Nussey, A. J. Wilson, and D. Réale. 2011. Measuring individual differences in reaction norms in field and experimental studies: A power analysis of random regression models. Methods Ecol. Evol. 2:362–374.

Mathot, K. J., P. J. van den Hout, T. Piersma, B. Kempenaers, D. Réale, and N. J. Dingemanse. 2011. Disentangling the roles of frequency-vs. state-dependence in generating individual differences in behavioural plasticity. Ecol. Lett. 14:1254–62.

McGuigan, K., A. Van Homrigh, and M. W. Blows. 2008. An evolutionary limit to male mating success. Evolution 62:1528–37.

Mitchell, D. J., and P. A. Biro. 2017. Is behavioural plasticity consistent across different environmental gradients and through time? Proc. R. Soc. B Biol. Sci. 284:20170893.

Mitchell, D. J., and T. M. Houslay. 2021. Context-dependent trait covariances: how plasticity shapes behavioral syndromes. Behav. Ecol. 32:25–29.

Moiron, M., K. J. Mathot, and N. J. Dingemanse. 2016. A multi-level approach to quantify speed-accuracy trade-offs in great tits (Parus major). Behav. Ecol. 27:1539–1546.

Morand-Ferron, J., E. Varennes, and L. A. Giraldeau. 2011. Individual differences in plasticity and sampling when playing behavioural games. Proc. R. Soc. B 278:1223–1230.

Niemelä, P. T., and N. J. Dingemanse. 2017. Individual versus pseudo-repeatability in behaviour: Lessons from translocation experiments in a wild insect. J. Anim. Ecol. 86:1033–1043.

Nussey, D. H., E. Postma, P. Gienapp, and M. E. Visser. 2005. Selection on heritable phenotypic plasticity in a wild bird population. Science 310:304–306.

Nussey, D. H., A. J. Wilson, and J. E. Brommer. 2007. The evolutionary ecology of individual phenotypic plasticity in wild populations. J. Evol. Biol. 20:831–844.

O’Dea, R. E., D. W. A. Noble, and S. Nakagawa. 2022. Unifying individual differences in personality, predictability and plasticity: A practical guide. Methods Ecol. Evol. 13:278–293.

Prestwich, K. N. 1994. The energetics of acoustic signaling in anurans and insects. Am. Zool. 34:625–643.

Price, T., and T. Langen. 1992. Evolution of correlated characters. Trends Ecol. Evol. 7:307–310.

R Development Core Team. 2021. R: A language and environment for statistical computing. R Foundation for Statistical Computing, Vienna, Austria.

Reichert, M. S. 2010. Aggressive thresholds in Dendropsophus ebraccatus: Habituation and sensitization to different call types. Behav. Ecol. Sociobiol. 64:529–539.

Reichert, M. S., and H. C. Gerhardt. 2012. Trade-offs and upper limits to signal performance during close-range vocal competition in gray tree frogs Hyla versicolor. Am. Nat. 180:425–437.

Reichert, M. S., and G. Höbel. 2018. Phenotypic integration and the evolution of signal repertoires: A case study of treefrog acoustic communication. Ecol. Evol. 8:3410–3429.

Reznick, D., L. Nunney, and A. Tessier. 2000. Big houses, big cars, superfleas and the costs of reproduction. Trends Ecol. Evol. 15:421–425.

Roff, D. 1997. Evolutionary Quantitative Genetics. Chapman and Hall, New York.

Roff, D. A., and D. J. Fairbairn. 2007. The evolution of trade-offs: where are we? J. Evol. Biol. 20:433–447.

Runkle, L. S., K. D. Wells, C. C. Robb, and S. L. Lance. 1994. Individual, nightly, and seasonal variation in calling behavior of the gray tree frog, Hyla versicolor: implications for energy expenditure. Behav. Ecol. 5:318–325.

Schwartz, J. J., B. Buchanan, and H. C. Gerhardt. 2002. Acoustic interactions among male gray treefrogs, Hyla versicolor, in a chorus setting. Behav. Ecol. Sociobiol. 53:9–19.

Schwartz, J. J., K. Huth, and T. Hutchin. 2004. How long do females really listen? Assessment time for female mate choice in the grey treefrog, Hyla versicolor. Anim. Behav. 68:533–540.

Sgrò, C. M., and A. A. Hoffmann. 2004. Genetic correlations, tradeoffs and environmental variation. Heredity 93:241–248.

Simmons, L. W., and D. J. Emlen. 2006. Evolutionary trade-off between weapons and testes. Proc. Natl. Acad. Sci. 103:16346–16351.

Sinervo, B., and E. Svensson. 2002. Correlational selection and the evolution of genomic architecture. Heredity 89:329–338.

Snell-Rood, E. C. 2013. An overview of the evolutionary causes and consequences of behavioural plasticity. Anim. Behav. 85:1004–1011.

Stamps, J. A. 2016. Individual differences in behavioural plasticities. Biol. Rev. 91:534–567.

Stamps, J., and P. Biro. 2016. Personality and individual differences in plasticity. Curr. Opin. Behav. Sci. 12:18–23.

Stearns, S. C. 1989. Trade-offs in life-history evolution. Funct. Ecol. 3:259–268.

Sueur, J., T. Aubin, and C. Simonis. 2008. Seewave, a free modular tool for sound analysis and synthesis. Bioacoustics 18:213–226.

Sullivan, B. K., and S. H. Hinshaw. 1992. Female choice and selection on male calling behaviour in the grey treefrog Hyla versicolor. Anim. Behav. 44:733–744.

Tanner, J. C., and M. A. Bee. 2020. Inconsistent sexual signaling degrades optimal mating decisions in animals. Sci. Adv. 6:eaax3957.

Tanner, J. C., J. L. Ward, R. G. Shaw, and M. A. Bee. 2017. Multivariate phenotypic selection on a complex sexual signal. Evolution 71:1742–1754.

van de Pol, M. 2012. Quantifying individual variation in reaction norms: how study design affects the accuracy, precision and power of random regression models. Methods Ecol. Evol. 3:268–280.

van Noordwijk, A. J., and G. de Jong. 1986. Acquisition and allocation of resources: Their influence on variation in life history tactics. Am. Nat. 128:137–142.

Walsh, B., and M. W. Blows. 2009. Abundant genetic variation + strong selection = multivariate genetic constraints: A geometric view of adaptation. Annu. Rev. Ecol. Evol. Syst. 40:41–59.

Ward, J. L., E. K. Love, A. Vélez, N. P. Buerkle, L. R. O. Bryan, and M. A. Bee. 2013. Multitasking males and multiplicative females: dynamic signalling and receiver preferences in Cope’s grey treefrog. Anim. Behav. 86:231–243.

Welch, A. M., M. J. Smith, and H. Carl Gerhardt. 2014. A multivariate analysis of genetic variation in the advertisement call of the gray treefrog, Hyla versicolor. Evolution 68:1629–1639.

Wells, K. D., and T. L. Taigen. 1986. The effect of social interactions on calling energetics in the gray treefrog (Hyla versicolor). Behav. Ecol. Sociobiol. 19:9–18.

Willi, Y., and J. Van Buskirk. 2022. A review on trade-offs at the warm and cold ends of geographical distributions. Philos. Trans. R. Soc. B Biol. Sci. 377:20210022.

Wolf, M., G. S. Van Doorn, and F. J. Weissing. 2011. On the coevolution of social responsiveness and behavioural consistency. Proc. R. Soc. B Biol. Sci. 278:440–448.

