## Supplementary Material for "Partitioning variance in a signaling trade-off under sexual selection reveals among-individual covariance in trait allocation"

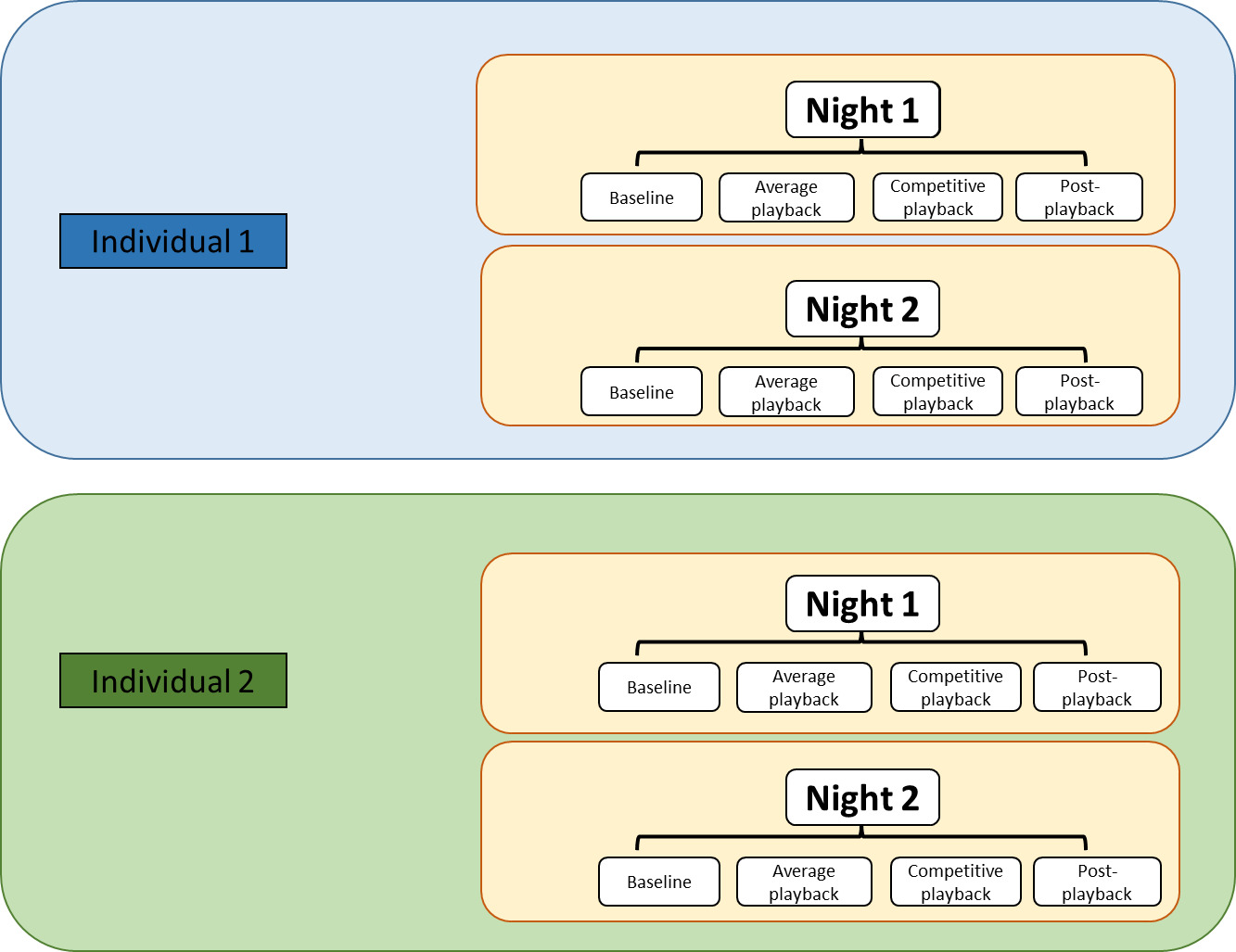

Figure S1. Schematic of the experimental design. On each night of recording for a given individual, we recorded its calling in four social contexts: baseline (i.e., spontaneous calling in its natural environment), in response to a playback of an average advertisement call, in response to a playback of a highly competitive advertisement call, and spontaneously calling after the playbacks. Whenever possible, we recorded each individual on multiple nights using the same procedure.

Table S1. Sources of phenotypic variance in inter-call interval across social contexts.

| Random effects | Context | Estimate [95% CI] |
| --- | --- | --- |
| Among-individual variance | Baseline | 0.0354 [0.0246, 0.0480] |
|  | Average | 0.0394 [0.0297, 0.0554] |
|  | Competitive | 0.0371 [0.0281, 0.0557] |
|  | Post-playback | 0.0490 [0.0356, 0.0688] |
| Within-individual variance | Baseline | 0.163 [0.156, 0.173] |
|  | Average | 0.141 [0.134, 0.146] |
|  | Competitive | 0.170 [0.164, 0.178] |
|  | Post-playback | 0.161 [0.152, 0.169] |
| Repeatability | Baseline | 0.177 [0.133, 0.231] |
|  | Average | 0.227 [0.173, 0.282] |
|  | Competitive | 0.193 [0.140, 0.247] |
|  | Post-playback | 0.253 [0.187, 0.307] |

Inter-call interval was calculated as the amount of time between the offset of one call and the onset of the next. Estimates show posterior modes with their associated 95% Credible Intervals for each variance component and the repeatability coefficient from the multivariate model of inter-call interval across social contexts. Note that inter-call interval was ln(x+1) transformed prior to analyses.

Table S2. Among-individual estimates for covariance and correlations in inter-call interval across social contexts.

| Contexts compared | Covariance [95% CI] | Correlation [95% CI] |
| --- | --- | --- |
| Average – baseline | 0.0307 [0.0200, 0.0405] | 0.808 [0.670, 0.877] |
| Competitive – baseline | 0.0283 [0.0188, 0.0385] | 0.756 [0.635, 0.870] |
| Competitive – average | 0.0325 [0.0237, 0.0459] | 0.844 [0.755, 0.925] |
| Post-playback – baseline | 0.0353 [0.0230, 0.0471] | 0.817 [0.718, 0.910] |
| Post-playback – average | 0.0359 [0.0240, 0.0488] | 0.758 [0.658, 0.876] |
| Post-playback – competitive | 0.0293 [0.0211, 0.0457] | 0.730 [0.592, 0.848] |

Estimates show the posterior mode and 95% Credible Intervals for the covariance and correlation at the among-individual level for inter-call interval across each pairwise combination of social contexts. Cross-context correlation coefficients that are clearly different from one give evidence for among-individual variance in inter-call interval plasticity, in other words that there is an individual-by-environment interaction. Comparing these results to those presented in Table 2 (showing values for call rate), it is noticeable that the correlation coefficients were very similar in all cases, justifying the use of call rate in our analyses.

Table S3. Within-individual estimates for covariance and correlations in inter-call interval across social contexts.

| Contexts compared | Covariance [95% CI] | Correlation [95% CI] |
| --- | --- | --- |
| Average – baseline | 0.0142 [−0.0590, 0.0749] | 0.0946 [−0.401, 0.479] |
| Competitive – baseline | −0.0410 [−0.0697, 0.0934] | −0.246 [−0.420, 0.550] |
| Competitive – average | −0.0263 [−0.0712, 0.0497] | −0.223 [−0.453, 0.328] |
| Post-playback – baseline | 0.00611 [−0.0820, 0.0511] | 0.0356 [−0.499, 0.315] |
| Post-playback – average | 0.0144 [−0.0566, 0.0702] | 0.140 [−0.374, 0.468] |
| Post-playback – competitive | 0.00229 [−0.0983, 0.0826] | 0.0125 [−0.608, 0.479] |

Estimates show the posterior mode with its associated 95% Credible Intervals for the covariance and correlation at the within-individual level for inter-call interval across each pairwise combination of social contexts. In all cases, estimates were very close to zero and credible intervals overlapped zero, indicating no evidence for significant within-individual covariance.

Table S4. Bivariate models testing for the covariance between call duration (CD) and inter-call interval (ICI)

|  | Average | Competitive |
| --- | --- | --- |
| Random effects | Estimate [95% CI] | Estimate [95% CI] |
| V_among-individual CD_ | 0.0133 [0.0106, 0.0184] | 0.0158 [0.0118, 0.0208] |
| V_among-individual ICI_ | 0.0405 [0.0294, 0.0556] | 0.0427 [0.0264, 0.0566] |
| COV_among-individual CD-ICI_ | 0.0153 [0.0106, 0.0225] | 0.0182 [0.0130, 0.0263] |
| *r*_among-individual CD-ICI_ | 0.695 [0.559, 0.791] | 0.791 [0.674, 0.875] |
| V_within-individual CD_ | 0.00573 [0.00549, 0.00596] | 0.00767 [0.00731, 0.00801] |
| V_within-individual ICI_ | 0.139 [0.135, 0.147] | 0.169 [0.163, 0.179] |
| COV_within-individual CD-ICI_ | 0.00100 [0.00028, 0.00188] | −0.00073 [−0.00185, 0.00041] |
| *r*_within-individual CD-ICI_ | 0.0289 [0.00896, 0.0658] | −0.0283 [−0.0501, 0.0126] |
| Phenotypic *r* (model) | 0.308 [0.216, 0.366] | 0.254 [0.193, 0.334] |
| Phenotypic *r* (raw) | 0.262 [0.216, 0.366] | 0.238 [0.210, 0.267] |

Random effect results from bivariate mixed-effects models of the correlations between call duration (CD) and inter-call interval (ICI) at the phenotypic, among-males, and within-males levels. Separate columns give results from separate models that were run on data from two different social contexts: the average and competitive playbacks. Estimates represent the posterior modes for the variance (V), covariance (COV) and correlation (*r*) estimates, with associated 95% Credible Intervals (CI). Note that inter-call interval was ln(x+1) transformed prior to analyses. The phenotypic *r* represents the posterior mode correlation coefficient calculated from the population data, either by calculating the correlation coefficient from the sum of all covariance estimates returned by the model, or by taking the correlation of the raw phenotypic values. Comparing these results to those presented in Table 3 in the main text (which shows the same analyses but using call rate instead of inter-call interval), it is noticeable that the results are very similar in all cases (except that the sign of the covariance is usually different because inter-call interval is related to the inverse of call rate), justifying the use of call rate in our analyses.

Table S5. Within-individual estimates for covariance and correlations in call characteristics across social contexts.

|  | Call duration |  | Call rate |  | Call effort |  |
| --- | --- | --- | --- | --- | --- | --- |
| Contexts compared | Covariance [95% CI] | Correlation [95% CI] | Covariance [95% CI] | Correlation [95% CI] | Covariance [95% CI] | Correlation [95% CI] |
| Average – baseline | 0.00003 [−5e−4, 0.00055] | 0.00469 [−0.0922, 0.104] | −0.00004 [−0.00035, 0.00038] | −0.0101 [−0.0815, 0.102] | 0.00002 [−0.00011, 0.00013] | 0.0118 [−0.0541, 0.0649] |
| Competitive – baseline | 0.00005 [0.00056, 0.00065] | 0.00905 [−0.0908, 0.103] | 0.00007 [−0.00029, 3e−4] | 0.0176 [−0.0792, 0.0815] | 0.00000 [−0.00011, 0.00013] | 0.00062 [−0.0527, 0.0674] |
| Competitive – average | 0.00005 [−0.00071, 0.00069] | −0.03453 [−0.110, 0.107] | −0.00006 [−0.00027, 0.00033] | −0.0170 [−0.0750, 0.0885] | −0.00002 [−0.00012, 0.00013] | −0.0122 [−0.0567, 0.0644] |
| Post-playback – baseline | −0.00001 [−0.00063, 0.00051] | −0.03732 [−0.111, 0.0899] | 0.00006 [−0.00031, 0.00029] | 0.0170 [−0.0857, 0.0819] | 0.00002 [−0.00011, 0.00012] | 0.0111 [−0.0574, 0.0639] |
| Post-playback – average | −0.00009 [−0.00068, 0.00062] | −0.0159 [−0.112, 0.108] | 0.00001 [−0.00026, 0.00031] | 0.00135 [−0.0711, 0.0849] | 0.00000 [−0.00011, 0.00012] | 0.00124 [−0.0562, 0.0629] |
| Post-playback − competitive | 0.00007 [−0.00074, 0.00071] | 0.0102 [−0.103, 0.107] | 0.00001 [−0.00026, 0.00025] | 0.00163 [−0.0779, 0.0787] | 0.00000 [−0.00012, 0.00011] | 0.00100 [−0.0614, 0.0554] |

Estimates show the posterior mode with its associated 95% Credible Intervals for the covariance and correlation at the within-individual level for each call characteristic across each pairwise combination of social contexts. In all cases, estimates were very close to zero and credible intervals overlapped zero, indicating no evidence for significant within-individual covariance.

Table S6. Repeatability coefficients from univariate models.

| Call characteristic | Context | Estimate [95% CI] |
| --- | --- | --- |
| Call duration | Baseline | 0.716 [0.656, 0.772] |
|  | Average playback | 0.716 [0.664, 0.775] |
|  | Competitive playback | 0.662 [0.617, 0.740] |
|  | Post-playback baseline | 0.631 [0.557, 0.686] |
|  | Cross-context | 0.597 [0.511, 0.646] |
| Call rate | Baseline | 0.224 [0.174, 0.285] |
|  | Average playback | 0.226 [0.170, 0.280] |
|  | Competitive playback | 0.174 [0.123, 0.219] |
|  | Post-playback baseline | 0.262 [0.207, 0.324] |
|  | Cross-context | 0.198 [0.158, 0.248] |
| Call effort | Baseline | 0.078 [0.052, 0.114] |
|  | Average playback | 0.103 [0.072, 0.140] |
|  | Competitive playback | 0.069 [0.041, 0.098] |
|  | Post-playback baseline | 0.109 [0.077, 0.154] |
|  | Cross-context | 0.114 [0.086, 0.143] |

Estimates show the posterior mode and 95% Credible Intervals for the repeatability estimate for each call characteristic in each social context. Repeatability coefficients were calculated from a univariate general linear mixed-effects model with the call characteristic of interest as the dependent variable, male identity as a random effect, and the same fixed effects as the multivariate model (see main text and Table S7). Compare these values obtained from the univariate model to the values reported from the multivariate model in Table 1. Cross-context refers to a single repeatability value that was calculated from data taken across all social contexts.

Table S7. Fixed effects from models of call characteristic plasticity.

|  | Call duration | Call rate | Call effort |
| --- | --- | --- | --- |
| Fixed effects | Estimate [95% CI] | Estimate [95% CI] | Estimate [95% CI] |
| Intercept (baseline) | 1.246 [1.205, 1.287] | 0.0646 [0.0334, 0.0954] | 0.154 [0.137, 0.172] |
| Intercept (average) | 1.407 [1.369, 1.449] | 0.0358 [0.00586, 0.0593] | 0.147 [0.130, 0.167] |
| Intercept (competitive) | 1.552 [1.502, 1.602] | 0.00058 [−0.0274, 0.0235] | 0.135 [0.111, 0.147] |
| Intercept (post-playback) | 1.283 [1.233, 1.3234] | 0.0131 [−0.0171, 0.0403] | 0.121 [0.0958, 0.136] |
| Temperature (baseline) | −0.0259 [−0.0279, −0.0248] | 0.00587 [0.00437, 0.00705] | −0.00122 [−0.00213, −0.00056] |
| Temperature (average) | −0.0312 [−0.0322, −0.0294] | 0.00622 [0.00472, 0.00702] | −0.00135 [−0.00217, −0.00056] |
| Temperature (competitive) | −0.0355 [−0.0370, −0.0329] | 0.00711 [0.00593, 0.00815] | −0.00049 [−0.0013, 0.00028] |
| Temperature (post-playback) | −0.0253 [−0.0272, −0.0235] | 0.00703 [0.00602, 0.00847] | 0.00003 [−0.00058, 0.00113] |
| Calendar day (baseline) | −0.00975 [−0.0154, −0.00567] | −0.0112 [−0.0151, −0.00765] | −0.00813 [−0.0106, −0.00604] |
| Calendar day (average) | −0.00574 [−0.0099, −0.00128] | −0.0122 [−0.0169, −0.0099] | −0.00848 [−0.0107, −0.00623] |
| Calendar day (competitive) | −0.00647 [−0.0112, 0.00042] | −0.0102 [−0.0130, −0.00705] | −0.00715 [−0.0095, −0.00532] |
| Calendar day (post-playback) | −0.00764 [−0.0143, −0.00391] | −0.00787 [−0.0117, −0.00477] | −0.00637 [−0.00834, −0.00346] |
| Time of night (baseline) | −0.00536 [−0.00906, −0.00182] | −0.00037 [−0.00294, 0.0028] | −0.00045 [−0.00253, 0.00096] |
| Time of night (average) | −0.0148 [−0.0175, −0.0107] | 0.00199 [−0.00115, 0.00457] | −0.00139 [−0.00335, 0.00021] |
| Time of night (competitive) | −0.0152 [−0.0184, −0.00992] | 0.00054 [−0.00258, 0.00264] | −0.00139 [−0.003, 0.00027] |
| Time of night (post-playback) | −0.0198 [−0.0228, −0.0149] | −0.00028 [−0.00275, 0.00298] | −0.00270 [−0.00419, −0.00057] |
| Year [2020] (baseline) | −0.135 [−0.149, −0.120] | 0.0211 [0.011, 0.0298] | −0.00624 [−0.0113, −0.00148] |
| Year [2020] (average) | −0.100 [−0.113, −0.0882] | 0.00849 [−0.00071, 0.0171] | −0.00583 [−0.0116, −0.00066] |
| Year [2020] (competitive) | −0.0919 [−0.107, −0.0756] | 0.00859 [0.00153, 0.0160] | −0.00393 [−0.00926, 0.00073] |
| Year [2020] (post-playback) | −0.0810 [−0.0975, −0.0655] | 0.0144 [0.00496, 0.0240] | −0.00578 [−0.0119, 0] |
| Sequence (baseline) | −0.00096 [−0.00134, −0.00056] | −0.00013 [−0.00035, 0.00039] | −0.00021 [−0.00044, 8e−05] |
| Sequence (average) | −0.00052 [−0.00066, −0.00024] | 0.00038 [2e−04, 6e−04] | 0.00020 [5e−05, 0.00034] |
| Sequence (competitive) | −0.00145 [−0.00178, −0.00119] | 0.00069 [0.00049, 0.00088] | 0.00021 [4e−05, 0.00035] |
| Sequence (post-playback) | −0.00300 [−0.00341, −0.00251] | 0.00039 [2e−05, 0.00074] | −0.00023 [−0.00049, 5e−05] |

Estimates show posterior modes and 95% Credible Intervals for fixed effects included in the models whose random effect variances are reported in Table 1 and covariances reported in Table 2. Each column corresponds to a multivariate model in which the specified call characteristic in each of the four social contexts was entered as the dependent variable, with fixed effects of temperature, calendar day, time of night, year and sequence. Temperature is the body temperature of the frog measured in °C. Calendar day is the day number of the year in which the recording was made, with 1 January being day 1. Time of night is in the units of hours past 9 PM. Both time of night and calendar day were standardized by centering and dividing by the standard deviation. Year is the year in which the recording was made, either 2019 (reference) or 2020. Sequence is a continuous variable numbering the calls in order on the recording to control for potential autocorrelation among the repeated observations within a recording.

Table S8. Fixed effects on the models of covariance between call duration and call rate.

|  | Average playback | Competitive playback |
| --- | --- | --- |
| Fixed effects | Estimate [95% CI] | Estimate [95% CI] |
| Intercept (call duration) | 1.425 [1.384, 1.465] | 1.529 [1.477, 1.582] |
| Intercept (call rate) | 0.0204 [−0.0182, 0.0428] | −0.00490 [−0.0326, 0.0241] |
| Call duration: Temperature | −0.0314 [−0.0329, −0.0299] | −0.0350 [−0.0368, −0.0325] |
| Call rate: Temperature | 0.00679 [0.00523, 0.00787] | 0.00722 [0.00595, 0.00853] |
| Call duration: Calendar day | −0.00034 [−0.00659, 0.00353] | −0.00862 [−0.0132, −5e−05] |
| Call rate: Calendar day | −0.0111 [−0.0154, −0.00812] | −0.00529 [−0.00904, −0.00261] |
| Call duration: Time of night | −0.0149 [−0.0189, −0.0114] | −0.0139 [−0.0184, −0.00885] |
| Call rate: Time of night | 0.00132 [−0.00137, 0.00446] | 0.00070 [−0.00192, 0.00369] |
| Call duration: Year [2020] | −0.1055 [−0.116, −0.0868] | −0.0659 [−0.0847, −0.0499] |
| Call rate: Year [2020] | 0.0139 [0.00457, 0.0240] | 0.0104 [0.00165, 0.0170] |
| Call duration: Sequence | −0.00043 [−0.00065, −2e−04] | −0.00135 [−0.00167, −0.00108] |
| Call rate: Sequence | 0.00049 [0.00022, 0.00063] | 0.00068 [0.00046, 0.00088] |

Variables as in Table S7. Columns correspond to separate models calculated for the average playback and the competitive playback. Separate effects are calculated for the two variables in these bivariate models, call duration and call rate.
